# Protein expression of prenyltransferase subunits in postmortem schizophrenia dorsolateral prefrontal cortex

**DOI:** 10.1101/654673

**Authors:** Anita L. Pinner, Toni M. Mueller, Khaled Alganem, Robert McCullumsmith, James H. Meador-Woodruff

## Abstract

The pathophysiology of schizophrenia includes altered neurotransmission, dysregulated intracellular signaling pathway activity, and abnormal dendritic morphology that contribute to deficits of synaptic plasticity in the disorder. These processes all require dynamic protein-protein interactions at cell membranes. Lipid modifications target proteins to membranes by increasing substrate hydrophobicity by the addition of a fatty acid or isoprenyl moiety, and recent evidence suggests that dysregulated post-translational lipid modifications may play a role in multiple neuropsychiatric disorders including schizophrenia. Consistent with these emerging findings, we have recently reported decreased protein S-palmitoylation in schizophrenia. Protein prenylation is a lipid modification that occurs upstream of S-palmitoylation on many protein substrates, facilitating membrane localization and activity of key intracellular signaling proteins. Accordingly, we hypothesized that in addition to palmitoylation, protein prenylation may be abnormal in schizophrenia. To test this, we assayed protein expression of the five prenyltransferase subunits (FNTA, FNTB, PGGT1B, RABGGTA, and RABGGTB) in postmortem dorsolateral prefrontal cortex from patients with schizophrenia and paired comparison subjects (N = 13 pairs). We found decreased levels of FNTA (14%), PGGT1B (13%), and RABGGTB (8%) in schizophrenia. To determine if upstream or downstream factors may be driving these changes, we also assayed protein expression of the isoprenoid synthases FDPS and GGPS1, and prenylation-dependent processing enzymes REC and ICMT. We found these upstream and downstream enzymes to have normal protein expression. To rule out effects from chronic antipsychotic treatment, we assayed FNTA, PGGT1B and RABGGTB in cortex from rats treated long-term with haloperidol decanoate, and found no change in the expression of these proteins. Given the role prenylation plays in localization of key signaling proteins found at the synapse, these data offer a potential mechanism underlying abnormal protein-protein interactions and protein localization in schizophrenia.

## Introduction

Altered neurotransmission is central to the pathophysiology of schizophrenia. Normal neurotransmission depends on regulation of receptor membrane localization and protein-protein interactions that regulate intracellular signaling activity.^1, 2^ Post-translational modifications (PTMs), including lipid modification of proteins, have been shown to regulate neuronal functions and intracellular pathways by facilitating dynamic protein-protein interactions at membranes.^3-5^ Altered post translational lipid modifications may mechanistically contribute to intracellular signaling abnormalities reported in schizophrenia.^6^

Post-translational lipid modifications include the enzymatic addition of an isoprenyl group such as farnesyl or geranylgeranyl (collectively called prenylation), or a fatty acid moiety, such as a palmitoyl or myristoyl group. Dysregulated lipid modifications of proteins have been implicated in neuropsychiatric disorders, including Alzheimer’s disease,^7^ Huntington’s disease,^8, 9^ and in a mouse model of schizophrenia.^10^ Abnormal lipid modifications have been reported in schizophrenia in dorsolateral prefrontal cortex (DLPFC), including decreased protein S-palmitoylation ^11^ and altered levels of a key N-myristoylated protein.^12^ While S-palmitoylation, N-myristoylation, and prenylation pathways can act independently, in some cases combinations of these lipid modifications are necessary for efficient membrane targeting, protein-protein interactions, and conformational dynamics of essential intracellular signaling proteins such as heterotrimeric G-proteins and small monomeric GTPases.^4, 5, 13-16^ Abnormal G-protein signaling has been implicated in schizophrenia,^6, 17-19^ and heterotrimeric G-protein subunits have been shown to require modifications by each one of these lipid modifications.^20-24^ Dysregulated prenylation could be a mechanism contributing to this illness via altered G-protein signaling.

Prenylation involves the addition of either farnesyl or geranylgeranyl isoprenoid group(s). Three prenylation enzymes are responsible for the thioether linkage of isoprenoid moieties—15-carbon farnesyl pyrophosphate (FPP) or 20-carbon geranylgeranyl pyrophosphate (GGPP)—to C-terminal cysteines: farnesyl transferase (FTase), geranylgeranyl transferase I (GGTase I), and geranylgeranyl transferase II (GGTase II).^25, 26^ Each enzyme is comprised of an α and β subunit. FTase and GGTase I have the same α subunit, FNTA, but have different β subunits: FNTB is the FTase β subunit, and PGGT1B the β subunit of GGTase I. GGTase II, which specifically geranylgeranylates Rab family proteins, is made of the RABGGTA and RABGGTB subunits.^26^ The addition of isoprenyl moieties leads to increased protein hydrophobicity, which facilitates targeted localization to membranes, lateral movement within membranes, and substrate conformational changes that can influence dynamic protein-protein interactions.^15, 16^

Given that prenylation occurs upstream of S-palmitoylation,^27^ which is decreased in schizophrenia,^11^ and facilitates the membrane targeting and resulting protein-protein interactions of key molecules associated with dynamic intracellular signaling,^4, 5, 28^ we hypothesized that farnesylation and/or geranylgeranylation is also dysregulated in schizophrenia. Our own bioinformatic analysis of publically available datasets reflects a pattern of transcript expression differences for prenylation-associated enzymes and substrates in schizophrenia that further suggests dysregulation of this pathway. Accordingly, we assayed protein expression of the prenyltransferases subunits FNTA, FNTB, PGGT1B, RABGGTA and RABGGTB, in DLPFC from schizophrenia and matched comparison subjects. To further characterize the regulation of this modification, we also assayed protein expression of the upstream isoprenoid synthases farnesyl diphosphate synthase (FDPS) and geranylgeranyl pyrophosphate synthase (GGPS1), and the prenylation-dependent downstream processing enzymes Ras-converting enzyme (RCE) and isoprenylcysteine carboxyl methyltransferase (ICMT) in the same subjects. To identify potential effects of antipsychotic treatment, we measured enzymes that were found altered in schizophrenia in cortex from rats chronically treated with haloperidol decanoate.

## Methods and Materials

### Human subjects

Samples of DLPFC (Brodmann Area (BA) 9/46) from schizophrenia and matched comparison subjects (N = 13 pairs) were obtained from the from the Mount Sinai School of Medicine (MSSM) NIH Brain and Tissue Repository (Table 1) as previously described.^12, 29^ Neuropathological examination of all subjects was conducted, and each subject’s medical history was reviewed extensively; detailed information regarding assessment is available http://icahn.mssm.edu/research/labs/neuropathology-and-brain-banking/neuropathology-evaluation. Subjects with previous drug or alcohol abuse, coma greater than 6 hours, suicide, or any evidence of neurodegenerative disease were excluded from the study. Next of kin consent was obtained for each subject. Subjects with schizophrenia all met DSM-III-R criteria, diagnosed by at least two clinicians, with documented onset of psychosis prior to 40 years of age, and a minimum of 10 years hospitalization for the illness. Comparison subjects were similarly evaluated and free of any neurological or psychiatric conditions. Based on our previous protein postmortem studies, power analysis determined this sample size was adequate to detect a moderate effect size ≥ 0.3 (α = 0.05, β = 0.2). We performed data analyses assuming equal variance as the subjects were matched pairs. Experimenters were blinded until data analyses.

**Table 1.** Paired Subject Demographics.

| Pair | Subject | Sex/Age | pH | PMI (hrs) |
| --- | --- | --- | --- | --- |
| 1 | Comparison | M/95 | 6.53 | 4.1 |
|  | Schizophrenia | M/97 | 6.5 | 9.3 |
| 2 | Comparison | F/66 | 6.85 | 22.6 |
|  | Schizophrenia | F/62 | 6.74 | 23.7 |
| 3 | Comparison | F/73 | 6.98 | 3 |
|  | Schizophrenia | F/70 | 6.51 | 13.2 |
| 4 | Comparison | M/70 | 6.1 | 6.7 |
|  | Schizophrenia | M/70 | 6.35 | 7.2 |
| 5 | Comparison | F/74 | 6.32 | 4.8 |
|  | Schizophrenia | F/75 | 6.49 | 21.5 |
| 6 | Comparison | M/73 | 6.17 | 14.9 |
|  | Schizophrenia | M/73 | 6.5 | 7.9 |
| 7 | Comparison | F/80 | 6.63 | 3.8 |
|  | Schizophrenia | F/81 | 6.67 | 15.1 |
| 8 | Comparison | F/79 | 6.38 | 5 |
|  | Schizophrenia | F/77 | 6.01 | 26.1 |
| 9 | Comparison | M/76 | 6.32 | 10.1 |
|  | Schizophrenia | M/80 | 6.37 | 9.7 |
| 10 | Comparison | F/85 | 7.27 | 2.9 |
|  | Schizophrenia | F/84 | 6.8 | 15.4 |
| 11 | Comparison | M/93 | 6.28 | 8 |
|  | Schizophrenia | M/92 | 6.67 | 21.9 |
| 12 | Comparison | F/89 | 6.72 | 4.2 |
|  | Schizophrenia | F/89 | 6.2 | 17.7 |
| 13 | Comparison | M/75 | 6.43 | 2.3 |
|  | Schizophrenia | M/78 | 6.64 | 9.6 |
| **Subject Summary** |  |  |  |  |
| Comparison (N=13) | | 79.1 ± 8.9 | 6.5 ± 0.3 | 7.1 ± 5.8 |
| Schizophrenia (N=13) | | 79.1 ± 9.7 | 6.5 ± 0.2 | 15.3 ± 6.5 |
| Abbreviations: postmortem interval (PMI), hours (hrs), female (F), male (M). |  |  |  |  |

### Antipsychotic Treated Rats

Animal studies and procedures were performed in accordance to institutional guidelines and approved by the Institutional Animal Care and Use Committee of the University of Alabama at Birmingham. Twenty male Sprague-Dawley rats (250g) were housed in pairs during the 9 month course of study. Haloperidol deconoate (28.5mg/kg, N=10) or vehicle (sesame oil, N=10) was administered every three weeks via intramuscular injection, for a total of 12 injections.^30, 31^ Animals were sacrificed by rapid decapitation, and the brains were harvested immediately. The right frontal cortex was dissected on wet ice, snap frozen in liquid nitrogen, and stored at −80°C. Sample sizes were determined by previous studies, experimenters were blinded until data analyses, and sample groups were randomized.

### Tissue Homogenization

Tissue samples from both human subjects and rats were homogenized in ice cold homogenization buffer (5mM Tris-HCl pH 7.5, 0.32M sucrose) supplemented with protease and phosphatase inhibitor tablets (Complete Mini, EDTA-free and PhosSTOP; Roche Diagnostics, Mannheim, Germany), using A Power Gen 125 (ThermoFisher Scientific, Rockford, Illinois) homogenizer at speed setting 5 for 60 seconds. A BCA protein assay kit (ThermoFisher Scientific) was used to determine protein concentration, and samples were stored at −80°C.

### Western Blot Analysis

Thawed homogenates were denatured at 70°C for 10min under reducing conditions, and stored at −20°C. Duplicate samples were loaded onto NuPAGE 4-12% Bis-Tris gels (Invitrogen, Carlsbad, CA) and transferred to nitrocellulose membranes using a BioRad Semi-Dry Transblotter (Hercules, CA). Membranes were incubated in Odyssey blocking buffer (LI-COR, Lincoln, NE) for 1 hr at room temperature (RT) before being probed with the primary antibody diluted in LI-COR blocking buffer with 0.1% Tween-20, using the conditions indicated in Table 2. After incubation in primary antibody, membranes were washed in cold Tris-buffered Saline + 0.05% Tween-20 (TBST) before being probed with IR-dye labeled secondary antibody diluted in LI-COR blocking buffer + 0.1% Tween-20 for 1 hr at RT. Finally, membranes were washed in cold TBST, then briefly rinsed in MilliQ water before being scanned with a LI-COR Odyssey imager. All antibodies were optimized for ideal conditions of each target protein within the linear range of detection for each assay, and ensuring the primary antibody was present in excess (Table 2). Valosin-containing protein (VCP) has been shown to be unchanged in multiple regions of schizophrenia brain^32, 33^ and was used as an intralane loading control for western blot normalization.

**Table 2.** Antibodies.

| Target Protein | Host Species | Dilution | Incubation | Company | Catalog # |
| --- | --- | --- | --- | --- | --- |
| FNTA | Rabbit | 1 : 1 000 | 16 hr $4^{\circ}\text{C}$ | Abcam | ab109738 |
| FNTB | Rabbit | 1 : 5 000 | 16 hr $4^{\circ}\text{C}$ | Abcam | ab109625 |
| RABGGTA | Rabbit | 1 : 500 | 16 hr $4^{\circ}\text{C}$ | Abcam | ab118781 |
| RABGGTB | Mouse | 1 : 1 000 | 16 hr $4^{\circ}\text{C}$ | Abnova | H00005876-M02 |
| PGGT1B | Mouse | 1 : 1 000 | 16 hr $4^{\circ}\text{C}$ | Abnova | H00005229-M02 |
| FDPS | Rabbit | 1 : 1 000 | 16 hr $4^{\circ}\text{C}$ | Abcam | ab153805 |
| GGPS1 | Rabbit | 1 : 1 000 | 16 hr $4^{\circ}\text{C}$ | Abcam | ab167168 |
| RCE1 | Rabbit | 1 : 500 | 16 hr $4^{\circ}\text{C}$ | Abcam | ab62531 |
| CMT | Rabbit | 1 : 500 | 16 hr $4^{\circ}\text{C}$ | Abcam | ab80872 |
| VCP | Mouse | 1 : 25 000 | 1 hr RT | Abcam | ab11433 |
| VCP | Rabbit | 1 : 25 000 | 1 hr RT | Abcam | ab109240 |

### Data Analysis

Protein expression was determined using LI-COR Odyssey 3.0 analytical software (Lincoln, NE). Intensity values were normalized to the intralane VCP intensity value after verifying that VCP was not changed between these schizophrenia and comparison subjects, consistent with previous reports.^32^ Duplicate values were averaged for each subject. All dependent measures were tested for normal distribution with the D’Agostino-Pearson omnibus normality test. Normally distributed data were analyzed using two-tailed paired Student’s t-tests, and Wilcoxon matched-pairs signed rank tests were used for non-normally distributed data using GraphPad Prism software (GraphPad Software, La Jolla, CA). No dependent measures were found to be associated with age, pH, or postmortem interval (PMI) using *post hoc* linear regression analyses. For all statistical tests, α = 0.05.

### Bioinformatic Analysis

We evaluated prenylation-associated targets for patterns of differential gene expression in schizophrenia using six publically available transcriptomic datasets generated from samples from the MSSM NIH Brain and Tissue Repository. These datasets include studies in gray matter homogenates from the middle temporal area (MTA; BA 21), temporopolar area (TPA; BA 38), anterior cingulate cortex (ACC; BA 32), and DLPFC (BA 46).^34^ Two datasets examined transcript levels in laser capture microdissected pyramidal neurons from superficial (lamina II-III) or deep (lamina V-VI) cortical layers of the DLPFC.^35^ The data were processed and analyzed using various R packages for differential expression analysis such as edgeR, DESeq2, and limma.^36-38^ The datasets are then aggregated and uniquely titled for processing. To visualize patterns of differential expression, a heatmap of log2 fold change values has been constructed to present harmonized data across all of the different datasets. Harmonization was achieved using empirical cumulative probabilities based on each dataset, and final harmonized values presented as standardized values that range from −1 to 1.^39^ The unsupervised clustering and construction of heatmaps were done using the pheatmap R package.^40^

## Results

### Prenyltransferase subunits are abnormally expressed in schizophrenia

We found that each prenyltransferase enzyme had decreased expression of either or both of its respective α and β subunits in schizophrenia relative to comparison subjects. The FTase and GGTase I α subunit, FNTA, was decreased 14% (*t*(12) = 3.74, p = 0.003). The GGTase I β subunit PGGT1B was decreased 13% (W = −77, p = 0.004) and the Rab protein-specific GGTase II β subunit RABGGTB was decreased 8% (*t*(12) = 2.29, p = 0.04) in schizophrenia (Figure 1). We also assayed FNTA, PGGT1B, and RABGGTB in rats chronically treated with haloperidol decanoate and found that haloperidol treatment did not affect expression of these proteins in these rats (Figure 2).

**Figure 1.**
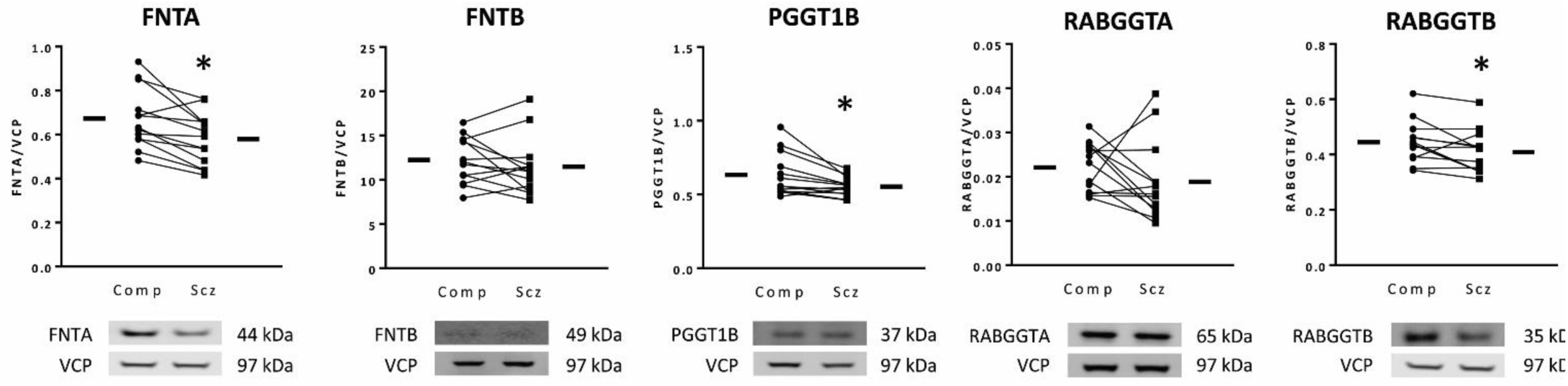
Decreased prenyltransferase subunit expression in schizophrenia. Western blots were used to assay protein expression in DLPFC of prenyltransferase subunits in schizophrenia (Scz) and matched, comparison subjects (Comp). FNTA, PGGT1B, and RABGGTB were decreased in schizophrenia. Data are expressed for each subject as the ratio of the signal intensity for protein of interest divided by the intensity of intralane VCP. Center lines are means. *p < 0.05

**Figure 2.**
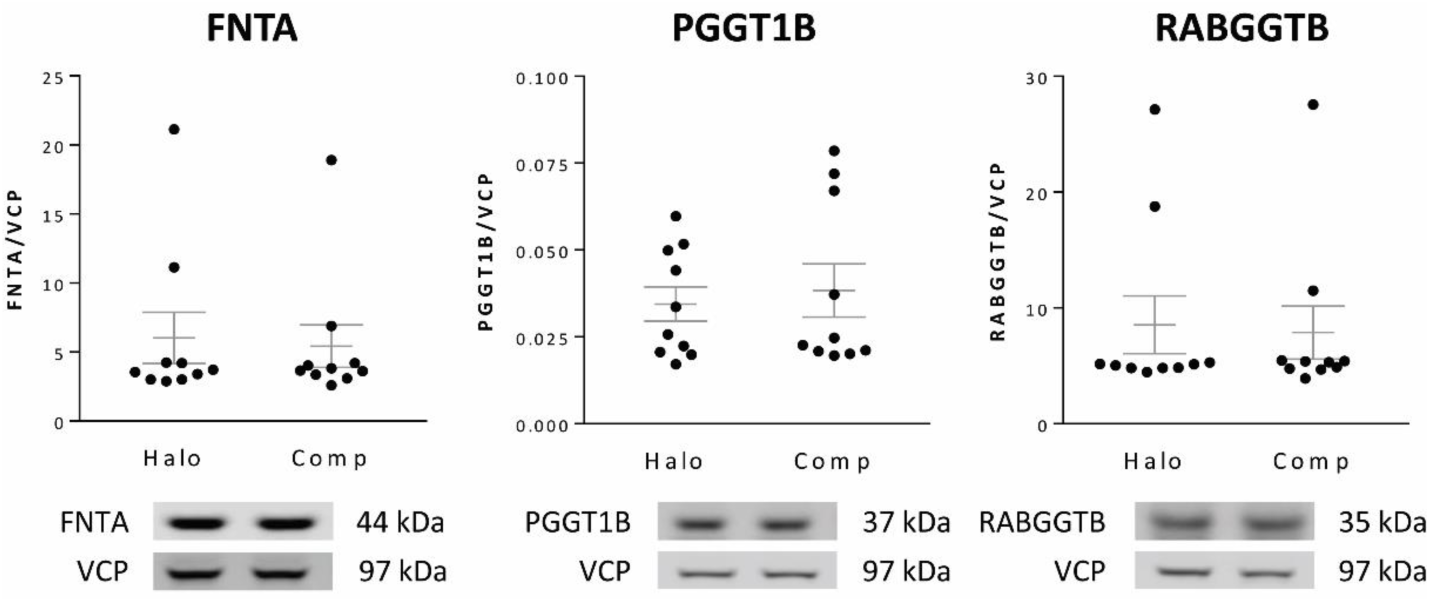
Chronic antipsychotic treatment does not alter expression of FNTA, PGGT1B and RABGGTB in rats. Western blotting techniques were used to assay protein expression of FNTA, PGGT1B, and RABGGTB in frontal cortex from adult male rats treated with haloperidol decanoate (28.5mg/kg every 3 weeks for 9 months; Halo) or vehicle (Comp). Data are expressed as a ratio of signal intensity for each target protein to intensity for VCP. Chronic haloperidol treatment did not change expression of these prenyltransferase subunits in rat cortex. Center lines are means ± S.E.M.

### Expression of upstream isoprenoid synthases and downstream prenylation-dependent enzymes is normal in schizophrenia

To determine if upstream lipid donor synthesis of FPP or GGPP was altered in schizophrenia, we assayed the synthases for these lipid-pyrophosphate molecules, farnesyl diphosphate synthase (FDPS) and geranylgeranyl pyrophosphate synthase (GPPS1). No differences in the expression of FDPS or GPPS1 were identified in schizophrenia (Table 3). To determine if enzymes downstream of prenyl attachment that are involved in the secondary processing of prenylproteins were altered, we measured protein expression of Ras-converting enzyme (RCE) and isoprenylcysteine carboxyl methyltransferase (ICMT) in schizophrenia and found normal expression of both (Table 3).

**Table 3.** Prenylation-associated enzyme expression levels in schizophrenia and comparison subjects.

|  | Comparison | Schizophrenia | Test statistic | p value |
| --- | --- | --- | --- | --- |
| **<u>Prenylsynthases</u>** |  |  |  |  |
| FDPS | 0.649 ± 0.067 | 0.641 ± 0.067 | *t* (12) = 0.35 | 0.73 |
| GGPS1 | 0.194 ± 0.066 | 0.189 ± 0.059 | *t* (12) = 0.36 | 0.72 |
| **<u>Prenyltransferase subunits</u>** |  |  |  |  |
| **FNTA** | **0.673 ± 0.136** | **0.580 ± 0.117** | ***t* (12) = 3.74** | **0.003\*** |
| FNTB | 12.24 ± 2.618 | 11.47 ± 3.235 | W = -25 | 0.41 |
| **PGGT1B** | **0.633 ± 0.146** | **0.553 ± 0.063** | **W = -77** | **0.004\*** |
| RABGGTA | 0.022 ± 0.006 | 0.019 ± 0.009 | *t* (12) = 1.14 | 0.28 |
| **RABGGTB** | **0.445 ± 0.076** | **0.409 ± 0.078** | ***t* (12) = 2.29** | **0.04\*** |
| **<u>Prenylcysteine processing enzymes</u>** |  |  |  |  |
| RCE | 0.127 ± 0.038 | 0.119 ± 0.033 | W = -11 | 0.74 |
| ICMT | 0.197 ± 0.061 | 0.188 ± 0.055 | *t* (12) = 0.68 | 0.51 |

Values are reported as means ± S.D.

### Transcript levels of prenylation-associated enzymes and prenylated substrates are altered in schizophrenia

Bioinformatic analysis of transcriptomic datasets generated from samples from the MSSM NIH Brain and Tissue Repository revealed that genes associated with prenylation demonstrate altered patterns of gene expression in schizophrenia. Genes encoding for upstream prenyl synthases, prenyltransferase subunits, prenylcysteine processing enzymes, and some GTPases (substrates of prenylation) exhibit differential expression relative to comparison subjects in one or more of the datasets evaluated (Supplementary Figure S1, Supplementary Table S1).

## Discussion

Neurotransmission, synaptic plasticity, dendritic dynamics, and protein subcellular localization have all been reported to be abnormal in schizophrenia. Prenylation is a cytosolic PTM that enables many GTPases associated with these processes to correctly localize for signaling transduction.^16, 19, 21, 26, 41-44^ GTPases have been shown to require combinations of lipid modifications including S-palmitoylation, N-myristoylation, and prenylation, which facilitate membrane-dependent GTPase activity.^20-24^ Given that a deficit in protein S-palmitoylation has been reported in schizophrenia,^45^ we hypothesized that abnormal prenylation may also contribute to altered G-protein signaling pathways implicated in the illness.^6, 17-19^ We found protein expression of FNTA, PGGT1B, and RABGGTB prenyltransferase subunits decreased in schizophrenia DLPFC relative to paired comparison subjects, changes not likely due to chronic antipsychotic treatment. Bioinformatic assessments identified patterns of differential gene expression of prenylation-associated enzymes and substrates in schizophrenia. For individual genes the direction and magnitude of differences appears to vary by brain region and cortical layer, however, identification of prenylation-associated differences across multiple datasets suggests that this functional pathway is involved in this illness. Together, these data are consistent with our previous findings of abnormal lipid modifications in schizophrenia, including abnormal S-palmitoylation and decreased expression of an N-myristoylated protein in schizophrenia DLPFC.^11, 12^

Given that FNTA, PGGT1B, and RABGGTB were decreased, each prenyltransferase enzyme αβ complex then has at least one abnormally expressed subunit, and GGTase I has reduced expression of both its α and β subunits. Previous reports demonstrated transcript-level upregulation of two of these subunits, FNTA and RABGGTB, in schizophrenia superior temporal gyrus (STG; Brodmann area 22)^46^ and prefrontal cortex (Brodmann areas 9 and 10),^47^ respectively. These changes reported for transcript expression are in the opposite direction of the protein expression changes identified in the current study, which might suggest that upstream or downstream regulatory molecules may also be altered in schizophrenia, or may reflect cellular compensation.

Since upstream or downstream factors could be driving changes in prenyltransferase expression, we also assayed protein expression of the isoprenoid synthases, FDPS and GGPS1, and prenylprotein processing enzymes, RCE and ICMT. FDPS and GGPS1 catalyze the production of the key intermediates in the mevalonate pathway that are the lipid donors for farnesylation and geranylgeranylation, FPP and GGPP, which are attached to proteins by their respective prenyltransferase(s).^26^ Following cytosolic prenylation, many prenylproteins are targeted to the endoplasmic reticulum (ER), where they are subject to additional processing steps before they can ultimately be trafficked to the correct membrane destination.^25^ For substrates of FTase and GGTase I that contain a C-terminal “CAAX” amino acid sequence (where C is cysteine, A is any aliphatic amino acid, and X is any amino acid), RCE cleaves the AAX residues from the prenyl-cysteine, and ICMT subsequently catalyzes the methylation of the prenyl-cysteine.^25^ These upstream and downstream enzymes, however, were not found to be differently expressed in schizophrenia DLPFC. Together, these data suggest that isoprenoid lipid donor synthesis and prenyl-cysteine modifications that occur following prenylation are normal in schizophrenia, but the actual attachment of prenyl groups to protein substrates could be impaired.

Many prenylproteins belong to the superfamily of G-proteins which play key roles in synaptic regulation. Heterotrimeric G-protein α, β, and γ subunits require lipid modifications for membrane targeting, protein interaction, and activity. Gα subunits can be N-myristoylated and/or S-palmitoylated,^14, 22, 48^ and Gγ subunits are prenylated.^16^ Prenylation is also required for some proteins involved with the regulation of Ca^2+^ signaling, spinogenesis, and synaptogenesis,^49, 50^ pathways which have been implicated in schizophrenia.^18, 43, 51, 52^

Many members of the Ras protein superfamily of small GTPases, which include Ras, Rho, Rab, Rap, Arf, Ran, and Rheb subfamilies, are also prenylated.^4^ The Rab and Rho protein subfamilies regulate membrane trafficking and cytoskeletal dynamics, as well as spatiotemporal aspects of vesicular recycling within the cell.^53^ These proteins require prenylation for correct interactions with their effector proteins at cell membranes.^54, 55^ Ras subfamily GTPases are involved in signal transduction and the regulation of gene expression^56^ and recent studies suggest Ras family protein signaling plays a role in memory formation.^57^ Given that pharmacological inhibition of Ras farnesylation prevents signal transduction to downstream targets in cell culture,^58^ decreased expression of functional FTases could inhibit the interaction of Ras proteins with their effectors and contribute to altered intracellular signaling and working memory in schizophrenia. Age-related downregulation of PGGT1B in mice is associated with decreased membrane-associated Rho-GTPases involved in synaptic plasticity. Inhibition of GGTase I *in vitro* leads to decreased protein levels of synaptophysin and GAP-43,^59^ and these proteins are also decreased in schizophrenia brain.^60-62^ Studies using statins, which inhibit the activity of the mevalonate pathway that produces FPP and GGPP, and prenylation-specific inhibitors in neuronal cultures, have reported decreased dendritic aborization^63, 64^ and increased axonal growth.^65, 66^ These reports emphasize that morphological abnormalities can arise from altered prenylation and suggest one potential mechanism contributing to dendritic spine abnormalities in schizophrenia brain.^43, 67^

Often the isoprenyl PTM by itself is not sufficient to maintain a strong membrane association, and many prenylated proteins require a second signal for membrane localization. This second signal is typically the addition of a palmitoyl group or the presence of a polybasic domain.^4, 68^ We have previously reported a deficit of protein S-palmitoylation in schizophrenia^11^ and it is likely that the subset of proteins which require both prenylation and S-palmitoylation are more susceptible to dysfunction if both of these pathways are impaired. Some members of the Ras subfamily are known to be both prenylated and S-palmitoylated, with prenylation occurring upstream of S-palmitoylation.^4, 27, 28^ Our prior report of decreased S-palmitoylation identified a widespread decrease of S-palmitoylation across many proteins, including Ras, but did not identify alterations in the expression levels of enzymes that attach or cleave palmitoyl groups that would explain the reduction.^11^ Our current finding of reduced prenyltransferase expression suggests that defective prenylation of Ras could potentially prevent the appropriate S-palmitoylation of these molecules, consistent with the findings of our previous study. If this is indeed the case, altered Ras lipidation might impair correct intracellular localization and/or signaling of these small GTPases. Deficits in DLPFC function, circuitry and working memory have been repeatedly implicated in schizophrenia.^69 70^ Considering Ras family signaling contributes to memory formation,^57^ abnormal lipid modifications in this brain region may contribute to memory-associated deficits in the disorder.

Postmortem brain studies in schizophrenia inherently have limitations. We evaluated gene expression differences and measured protein expression levels in aged subjects, and these data may not generalize to different age groups or earlier stages of the illness. Our protein study is also restricted to the DLPFC, thus additional brain regions will need to be examined to determine if these findings are widespread or brain region-specific. Furthermore, given that some enzymes measured in this study demonstrate cell type-specific patterns of expression, prenylation pathway alterations identified here may preferentially impact a distinct subpopulation of cells and cell-type specific protein measures will be necessary to investigate this possibility. Another major limitation of schizophrenia postmortem studies is that chronic antipsychotic treatment may affect expression of some proteins independent of the disorder. To rule out potential effects of long-term antipsychotic use, we assayed these proteins in frontal cortex of rats chronically treated with haloperidol decanoate. We did not find any effects of long-term haloperidol treatment on protein levels of FNTA, PGGT1B, or RABGGTB, which suggests that reduced prenyltransferase subunit expression identified in schizophrenia is likely due to the illness and not a result of chronic antipsychotic treatment. However, it is important to note that the haloperidol studies were not performed in an animal model of schizophrenia, and potential interactions between disease pathophysiology and long-term antipsychotic use that may influence protein prenylation cannot be completely ruled out.

In summary, we found decreased protein expression of the prenyltransferase subunits FNTA, PGGT1B, and RABGGTB in schizophrenia DLPFC as well as evidence from a bioinformatic analysis of differential gene expression of prenylation-associated genes across multiple brain regions and cortical layers. These data are consistent with other evidence of abnormal lipid modifications in schizophrenia^11, 12^ and suggest a potential mechanism for our report of reduced Ras S-palmitoylation in the face of normal palmitoyltransferase expression. Because of its importance in G protein signaling and small GTPase activity, abnormal prenylation is also a potential mechanism underlying altered intracellular signaling, dendritic dynamics, and subcellular protein localization previously reported in schizophrenia. Decreased protein expression of these prenyltransferase subunits could contribute to many facets of the pathophysiology of schizophrenia by its important role in multiple cell biological processes.

## Acknowledgements

The authors gratefully acknowledge Dr. Rosalinda Roberts and the Alabama Brain Collection for providing materials used to optimize conditions used in this study.

## Conflict of Interest Statement

All authors declare that they have no conflicts of interest.

