## Supplementary Figure S1 for "Protein expression of prenyltransferase subunits in postmortem schizophrenia dorsolateral prefrontal cortex"

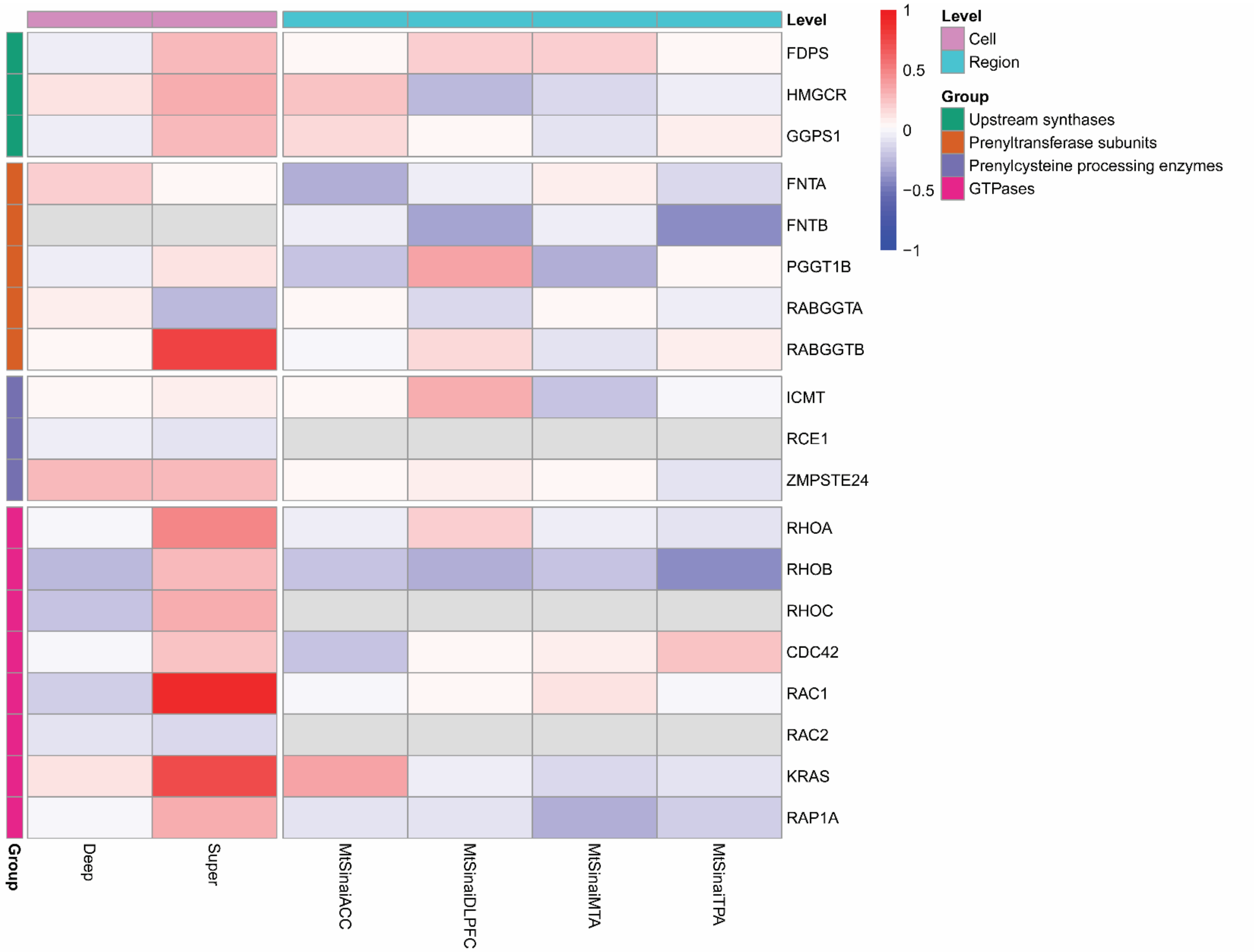


**Supplementary Figure S1. Transcript expression differences of prenylation-associated genes in multiple brain regions in schizophrenia.** Heatmap of log2 fold change values harmonized across datasets. Fold change, log2 fold change, and p-values are reported in Supplementary Table S1. Genes with increased expression relative to comparison subjects are indicated in shades of red, while genes with decreased expression are indicated in shades of blue. Gray indicates that the gene was not measured in the corresponding dataset. Transcriptomic datasets from schizophrenia and comparison subjects included in the heatmap were generated from tissue samples from the MSSM NIH Brain and Tissue Repository from the following brain regions: middle temporal area (MtSinaiMTA), temporopolar area (MtSinaiTPA), anterior cingulate cortex (MtSinaiACC), and dorsolateral prefrontal cortex (MtSinaiDLPFC). Transcriptomic datasets generated from laser capture microdissected pyramidal neurons from superficial (lamina II-III; Super) or deep (lamina V-VI; Deep) cortical layers of DLPFC are also included. Genes listed on the y-axis are grouped by function; on the x-axis, datasets are grouped by the specificity level of the study from which they were obtained (Cell or Region).
