## Supplementary Table S1 for "Protein expression of prenyltransferase subunits in postmortem schizophrenia dorsolateral prefrontal cortex"

**Supplementary Table S1. Expression differences of prenylation-associated genes from transcriptomic datasets**

| | | MtSinaiMTA | | | MtSinaiTPA | | | MtSinaiACC | | | MtSinaiDLPFC | | | DLPFC_Super | | | DLPFC_Deep | | |
|---|---|---|---|---|---|---|---|---|---|---|---|---|---|---|---|---|---|---|---|
| | Gene Symbol | FC | LogFC | p-value | FC | LogFC | p-value | FC | LogFC | p-value | FC | LogFC | p-value | FC | LogFC | p-value | FC | LogFC | p-value |
| Upstream synthases | FDPS | **1.15** | **0.20** | 0.05 | 1.02 | 0.02 | 0.81 | 1.04 | 0.06 | 0.62 | **1.16** | **0.21** | 0.28 | **1.22** | **0.28** | 0.19 | -1.02 | -0.03 | 0.74 |
| | HMGCR | **-1.10** | **-0.14** | 0.34 | -1.04 | -0.05 | 0.81 | **1.19** | **0.26** | 0.23 | **-1.18** | **-0.24** | 0.15 | **1.24** | **0.31** | 0.22 | 1.09 | **0.13** | 0.45 |
| | GGPS1 | -1.06 | -0.09 | 0.54 | 1.06 | 0.09 | 0.44 | **1.14** | **0.18** | 0.13 | 1.03 | 0.05 | 0.66 | **1.21** | **0.28** | 0.19 | -1.04 | -0.05 | 0.60 |
| Prenyltransferase subunits | FNTA | 1.06 | 0.08 | 0.67 | -1.08 | -0.11 | 0.25 | **-1.21** | **-0.27** | 0.07 | -1.01 | -0.02 | 0.90 | 1.04 | 0.05 | 0.74 | **1.17** | **0.22** | 0.06 |
| | FNTB | -1.03 | -0.04 | 0.87 | **-1.34** | **-0.42** | **0.004** | -1.03 | -0.05 | 0.77 | **-1.25** | **-0.32** | 0.08 | | | | | | |
| | PGGT1B | **-1.21** | **-0.28** | 0.09 | 1.03 | 0.04 | 0.79 | **-1.15** | **-0.20** | 0.33 | **1.31** | **0.39** | 0.21 | **1.10** | **0.14** | 0.49 | -1.04 | -0.06 | 0.55 |
| | RABGGTA | 1.03 | 0.04 | 0.73 | -1.03 | -0.05 | 0.71 | 1.02 | 0.02 | 0.88 | **-1.10** | **-0.14** | 0.19 | **-1.17** | **-0.23** | 0.05 | 1.05 | 0.07 | 0.47 |
| | RABGGTB | -1.06 | -0.09 | 0.50 | 1.05 | 0.06 | 0.57 | -1.00 | -0.01 | 0.96 | **1.11** | **0.15** | 0.39 | **1.71** | **0.78** | 0.08 | 1.02 | 0.03 | 0.85 |
| Prenylcysteine processing enzymes | ICMT | **-1.17** | **-0.22** | **0.01** | 1.01 | 0.01 | 0.94 | 1.04 | 0.06 | 0.69 | **1.25** | **0.32** | 0.41 | 1.06 | 0.09 | 0.42 | 1.02 | 0.03 | 0.74 |
| | RCE1 | | | | | | | | | | | | | -1.05 | -0.07 | 0.52 | -1.02 | -0.02 | 0.80 |
| | ZMPSTE24 | 1.03 | 0.04 | 0.65 | -1.06 | -0.09 | 0.16 | 1.04 | 0.06 | 0.58 | 1.07 | 0.09 | 0.35 | **1.20** | **0.27** | 0.17 | **1.20** | **0.27** | 0.09 |
| GTPases | RHOA | -1.02 | -0.03 | 0.77 | -1.05 | -0.07 | 0.38 | -1.02 | -0.03 | 0.80 | **1.15** | **0.20** | 0.33 | **1.41** | **0.49** | **0.04** | -1.01 | -0.02 | 0.89 |
| | RHOB | **-1.15** | **-0.21** | 0.15 | **-1.33** | **-0.41** | **0.002** | **-1.14** | **-0.19** | 0.18 | **-1.21** | **-0.28** | 0.10 | **1.22** | **0.29** | **0.02** | **-1.18** | **-0.23** | **0.01** |
| | RHOC | | | | | | | | | | | | | **1.24** | **0.31** | **0.03** | **-1.15** | **-0.21** | 0.08 |
| | CDC42 | 1.05 | 0.06 | 0.69 | **1.18** | **0.23** | **0.03** | **-1.16** | **-0.22** | 0.29 | 1.03 | 0.04 | 0.76 | **1.19** | **0.25** | 0.29 | 1.01 | 0.01 | 0.92 |
| | RAC1 | **1.10** | **0.13** | 0.07 | 1.00 | 0.00 | 0.95 | 1.01 | 0.02 | 0.82 | 1.02 | 0.03 | 0.66 | **1.84** | **0.88** | 0.06 | **-1.11** | **-0.15** | 0.32 |
| | RAC2 | | | | | | | | | | | | | **-1.10** | **-0.13** | 0.21 | -1.06 | -0.09 | 0.51 |
| | KRAS | **-1.10** | **-0.14** | 0.25 | -1.05 | -0.07 | 0.58 | **1.31** | **0.38** | **0.01** | -1.04 | -0.05 | 0.61 | **1.65** | **0.73** | **0.01** | 1.09 | **0.13** | 0.38 |
| | RAP1A | **-1.22** | **-0.29** | 0.06 | **-1.13** | **-0.18** | 0.16 | -1.04 | -0.06 | 0.73 | -1.04 | -0.06 | 0.75 | **1.24** | **0.32** | 0.16 | 1.01 | 0.01 | 0.91 |

**Values in bold type indicate fold change (FC) ≥ ± 1.10 or $p < 0.05$**
